# Hydra domain drives SNF2L multimerization and marks ISWI diversification in parasites

**DOI:** 10.1101/2025.09.03.673926

**Authors:** Belen Pachano, Jill von Velsen, Charlotte Corrao, Caroline Mas, Léa Pounot, Mohamed-Ali Hakimi, Matthew W. Bowler, Christopher Swale

## Abstract

ISWI chromatin remodelers are conserved regulators of nucleosome positioning and chromatin accessibility across eukaryotes, yet their evolutionary diversification is poorly understood. In the apicomplexan parasite *Toxoplasma gondii*, we identify Hydra, a previously unrecognized globular domain embedded within *Tg*SNF2L, one of two ISWI paralogues. Hydra is structurally unique, lacking homology to any known protein fold, and represents a lineage-specific insertion in an otherwise structurally conserved protein family. Biochemical analyses reveal that the isolated Hydra domain self-assembles into stable oligomers, undergoing reversible equilibrium with its monomeric form. Cryo-electron microscopy analysis reveals discrete globular assemblies, though little consistency could be obtained, suggesting a highly dynamic complex. Deletion of Hydra from full-length *Tg*SNF2L disrupts its intrinsic ability to form higher order oligomers in solution, yielding predominantly monomeric and dimeric species. Functionally, the Hydra-driven multimerization of *Tg*SNF2L modulates its availability for chromatin engagement in response to cell-cycle cues. Hydra thus represents the first reported structural innovation within the ISWI family.

## Introduction

In eukaryotic cells, DNA is tightly wrapped around an octamer of histones to form nucleosomes. Despite their stability, nucleosomes are inherently dynamic, a property largely dependent on the activity of ATP-dependent chromatin remodelers. These molecular machines generate the force required to disrupt histone-DNA contacts, enabling selective access to regulatory DNA elements. In densely packed chromatin, critical cis-regulatory sites can remain inaccessible, since DNA-binding proteins cannot engage sequences positioned within the nucleosome core. Consequently, the packaging of DNA into chromatin provides a fundamental layer of genomic regulation by modulating the accessibility of key regulatory elements to proteins such as transcription factors and polymerase initiation complexes, thereby facilitating essential biological processes such as transcription initiation, elongation and termination, DNA replication and DNA repair, all of which rely on dynamic chromatin rearrangements.

Chromatin remodelers are classified into distinct well-defined families, including SWI/SNF, INO80, ISWI, and CHD, distinguished by the characteristics of their Snf2-type motor ATPases (Flaus et al. 2006; Zhou et al. 2016). These motors have been shown to carry out most of the biochemical activities of their parent complexes by directionally translocating DNA using chemomechanical cycles of ATP binding and hydrolysis. On nucleosomes, this translocation alters and disrupts histone–DNA contacts, catalyzing DNA mobilization around histones (‘nucleosome sliding’), histone ejection, and the exchange of histone variants or (un)modified histones. However, despite the shared sequence homology within their ATPase domains, SWI/SNF, INO80, ISWI, and CHD classes play distinct roles *in vivo* and have distinct biochemical behaviors (Clapier and Cairns 2009; Narlikar et al. 2013; Zhou et al. 2016). SWI/SNF can reposition or completely eject nucleosomes (nucleosome sliding and disassembly, respectively), whereas ISWI and CHD primarily slide nucleosomes that are affected by extra-nucleosomal DNA flanking a nucleosome and the histone H4 tail (Narlikar et al. 2013). Remarkably, the human ISWI remodeler SNF2h is most effective as a dimer, whereas SWI/SNF and CHD function mainly as monomeric ATPases (Eustermann et al. 2024).

In addition to the helicase domain, chromatin remodelers also possess additional accessory domains that are crucial for their specialized functions. For example, SWI/SNF proteins commonly feature bromodomains that recognize acetylated histones (Filippakopoulos and Knapp 2014), while members of the CHD family typically include chromodomains that play a role in chromatin targeting and protein-protein interactions (Alendar and Berns 2021). The ISWI family is typified by the occurrence of a C-terminal domain termed HAND SANT-SLIDE (HSS). Within the HSS, the SANT domain recognizes nucleosomal surfaces, whereas the SLIDE domain interacts with linker DNA, enabling ISWI to sense nucleosome spacing (Grüne et al. 2003; Dang and Bartholomew 2007; Zhou et al. 2016). By guiding the ATPase motor’s engagement with nucleosomal DNA, these domains collectively modulate both the efficiency and the directionality of nucleosome repositioning. While other chromatin remodeler families (such as CHD) frequently incorporate diverse chromatin-recognition modules to guide their genomic targeting, ISWI remodelers rely exclusively on domains that facilitate their ATP-dependent sliding activity. Given this functional constraint, the domain architecture of ISWI proteins has remained largely unaltered throughout evolution.

The Apicomplexa encompass a large and diverse phylum of obligate intracellular protozoan parasites, including the causative agents of malaria, toxoplasmosis, and cryptosporidiosis. Their evolutionary history features an endosymbiotic event between a red alga and a heterotrophic eukaryotic protist, and they constitute one of the few entirely parasitic eukaryotic lineages. The transition to parasitism was marked by extensive gene loss, particularly of those genes associated with free-living lifestyles, as well as lineage-specific expansions of factors essential for host invasion, immune evasion, and intracellular survival (White and Suvorova 2018). This specific evolutive route has led apicomplexan parasites to possess chimeric genomes combining plant-like genes inherited from their algal endosymbiont, metazoan-derived genes laterally transferred from animal hosts, and “newly” evolved sequences, together reflecting the specialized adaptations of these parasites.

We recently showed that *Toxoplasma gondii* encodes two unique ISWI remodelers, *Tg*SNF2h and *Tg*SNF2L, with distinct features and functions (Pachano et al. 2025). *Tg*SNF2h assembles into a complex that actively modulates chromatin accessibility to regulate stage-specific gene expression, whereas *Tg*SNF2L associates with AP2X-4 and RSF1-like proteins to form two separate complexes not linked to transcriptional control. Sequence based and structural predictions using AlphaFold-3 (Abramson et al. 2024) indicate that both *Tg*SNF2h and *Tg*SNF2L possess ATPase and HSS domains closely mirroring those of their human counterparts. Notably, *Tg*SNF2L contains an additional “middle” domain, previously termed “domain of unknown function” (DUF), absent from all other ISWI-related proteins, including *Tg*SNF2h, for which no structural homologs were identified in publicly available AlphaFold/UniProt databases using Foldseek structural searches (van Kempen et al. 2024). The emergence of a novel structural domain within the ISWI family of remodelers, as revealed in *Tg*SNF2L, represents an unprecedented finding. Here, we provide structural and functional characterization of this DUF, which we refer to as the “Hydra domain”, a novel coccidia-specific insertion that serves as a multimerization module in *Tg*SNF2L and constitutes the first known acquisition of a novel structural domain in this otherwise highly conserved remodeler family.

## Results

### Identification of a novel evolutionary hallmark of coccidian ISWI proteins

Snf2-type ATPases are defined by a conserved helicase module composed of two RecA-like folds, often referred to as the DEXDc and HELICc domains, that couple ATP binding and hydrolysis to DNA translocation (Narlikar et al. 2013; Zhou et al. 2016; Eustermann et al. 2024). In an earlier study, we showed that apicomplexan parasites encode the four canonical Snf2-type remodeler families, including ISWI (Pachano et al. 2025). Building on those findings, we now performed a phylogenetic analysis of full-length ISWI proteins from *Homo sapiens, Saccharomyces cerevisiae, Arabidopsis thaliana*, and multiple apicomplexan species. We observed that apicomplexan ISWI homologs segregate into two distinct clades, a *Tg*SNF2h-like group and a *Tg*SNF2L-like group, diverging from non-parasitic counterparts (**Fig. 1a**). Notably, some apicomplexans retain both homologs, whereas others have selectively lost one, suggesting lineage-specific retention patterns and potential functional divergence.

**Fig. 1.**
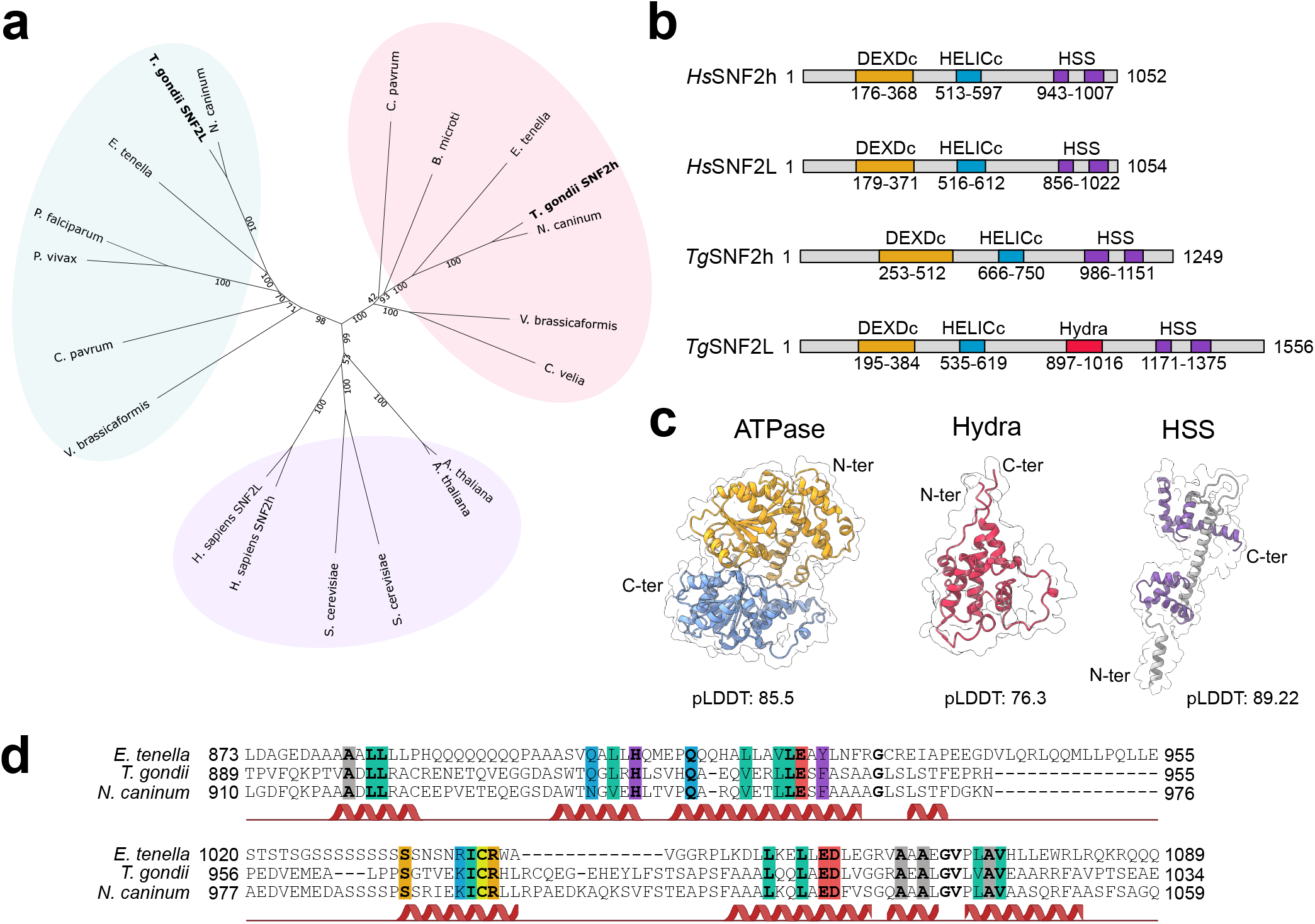
A novel coccidian-specific domain, Hydra, defines a unique ISWI domain architecture in *Toxoplasma gondii*. **a**. Neighbor-joining radial phylogenetic analysis of full-length ISWI proteins from apicomplexan parasites (*T. gondii, N. caninum, E. tenella, P. falciparum, P. vivax, B. microti, C. parvum*), chromerids (*C. velia, V. brassicaformis*), and model eukaryotes (*A. thaliana, H. sapiens, S. cerevisiae*) reveals two distinct apicomplexan-specific clades corresponding to *Tg*SNF2h- and *Tg*SNF2L-like proteins. **b**. Schematic domain organization of human and *T. gondii* ISWI proteins highlights the presence of a unique insertion in *Tg*SNF2L, termed Hydra. **c**. Secondary structures and surface of the domains of *Tg*SNF2L predicted by AlphaFold v2; C-ter; C-terminal; N-ter, N-terminal. **d**. Multiple sequence alignment of the Hydra domain from coccidian SNF2L-like ISWI proteins showing conserved residues and predicted secondary structures.

Phylogenetic analyses indicate that the last common ancestor of apicomplexan parasites harbored two ISWI paralogs, a SNF2h-like and a SNF2L-like remodeler, as evidenced by their presence in *Vitrella brassicaformis*, a free-living relative of apicomplexans (**Fig. 1a**). Coccidian parasites such as *T. gondii* have retained both homologs, exhibiting greater divergence than that observed in mammals. Notably, *Tg*SNF2L features a previously uncharacterized coccidia-specific insertion absent in ISWI proteins from model organisms (**Fig. 1b**), as well as from *Plasmodium* and *Cryptosporidium* species. We termed this insertion the Hydra domain (previously referred to as a DUF) (Pachano et al. 2025), which AlphaFold3 (Abramson et al. 2024) predicts to form five α-helices with good confidence (average pLDDT >70, albeit supported by few aligned sequences). Also present in *Neospora caninum* and *Eimeria tenella*, this domain appears restricted to the coccidian subgroup, underscoring its unique evolutionary origin. Sequence conservation within Hydra is substantially lower than in the ATPase or HSS domains, with variability concentrated in predicted loop regions and conserved residues primarily located within predicted α-helices (**Fig. 1d**).

### Recombinant Hydra domain self-assembles into a reversible decamer

To investigate the potential function of the Hydra domain, we recombinantly expressed the small domain in *E. coli* with a C-terminal histidine tag. The protein is abundantly produced and can be easily purified after a Ni-NTA pulldown. However, size exclusion chromatography (SEC) revealed two well-separated elution peaks (**Fig. 2a, b**). SEC coupled to multi-angle laser light scattering (SEC-MALLS, **Fig. 2c**) showed that the later-eluting peak corresponds to a monomer with an estimated mass of 18.5 kDa (theoretical mass of 17.8 kDa), whereas the earlier peak corresponds to a decamer with an estimated molar mass of 181.1 kDa. Interestingly, the oligomeric distribution is a reversible dynamic equilibrium, as the re-injection of purified decamer or monomer fractions reproducibly regenerated both species (**Fig. 2b, Supp Fig. 1A**). Native gel electrophoresis of SEC-separated complexes identified only 3 main species (**Supp Fig. 1A-C**): the monomer (migrating as a double band), the decamer, and a minor higher molecular weight species consistent with a putative 20-mer. Mass photometry of a 200 nM protein solution detected a large multimer with an estimated mass of ∼180 kDa (**Fig. 2d**), whereas the monomer was below the instrument’s 30 kDa detection limit, indicating that the high-molecular-weight decamer persists even at low protein concentrations.

**Fig. 2.**
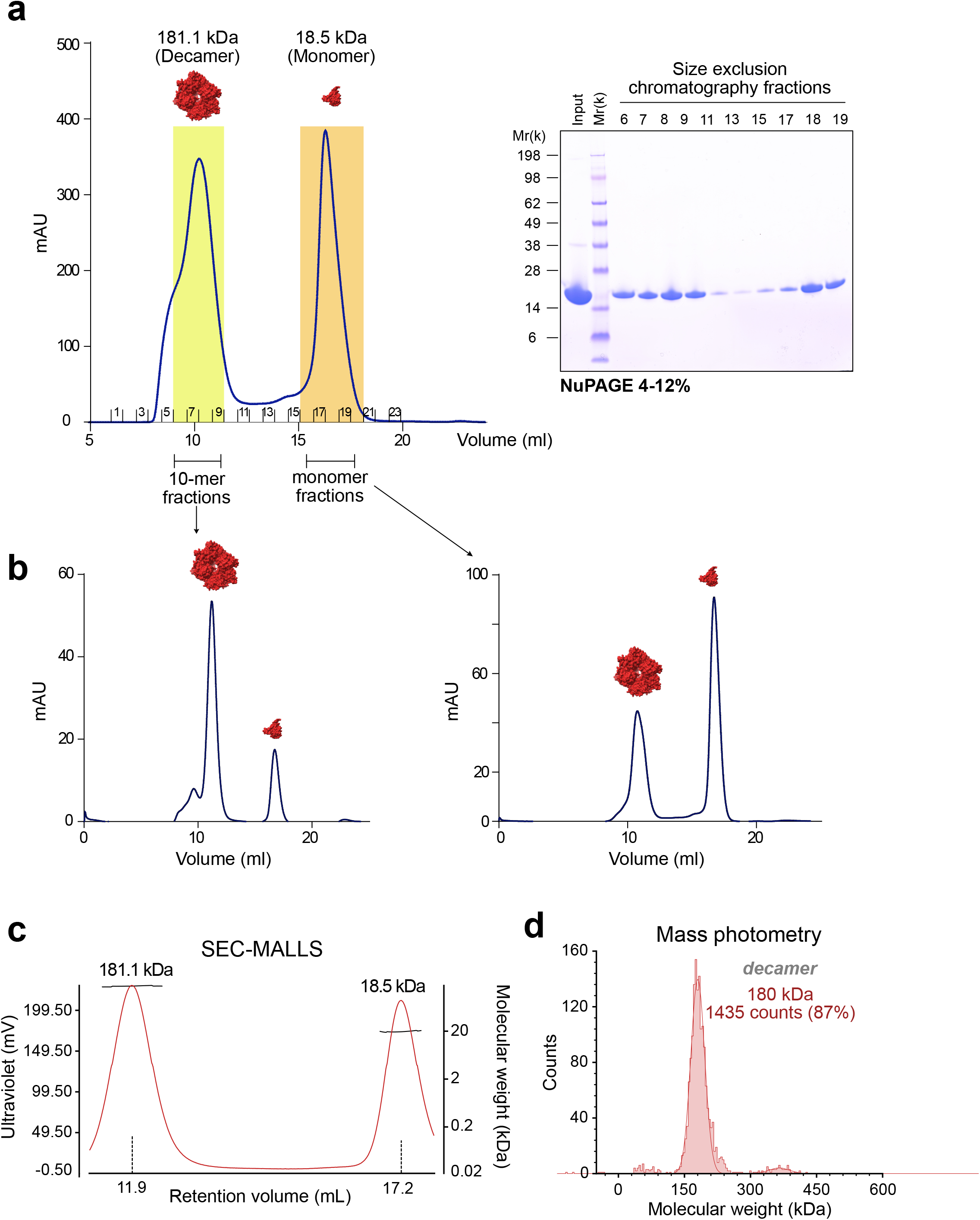
*Tg*SNF2L middle domain, Hydra, oligomerizes in a reversible and dynamic equilibrium *in vitro*. **a**. Size exclusion chromatography using a S200 10/300 increase and corresponding NuPAGE gel of the eluted fractions performed on Ni-NTA purified Hydra reveals two distinct peaks. The first peak corresponds to a decamer in solution as revealed by SEC MALLS (**c, d**) and second peak corresponds to a monomer. **b**. Upon re-injection on the same S200 10/300 increase column of either decamer or monomer peak fractions, two monomer/decamer peaks dynamically form again. SEC MALLS and mass photometry measurements both confirm that Hydra forms either an 18 kDa monomer and a 180 kDa decamer. **a**. SEC MALLS chromatogram of re-injected hydra monomer fractions onto an S200 10/300 increase column displays two distinct UV 280 nm peaks with the corresponding molecular weight estimates using MALLS. **b**. Mass photometry accurately detects only the decamer of Hydra as the monomeric form is well below the detection range of the MP one (40 kDa), higher oligomeric forms are also partially detected using this method.

Using negative-stain transmission electron microscopy (EM), we analyzed the decamer peak fractions from gel filtration and observed homogeneous, globular particles (**Fig. 3a**), suggesting a specific assembly fold. Larger aggregates were also visible, indicating the potential for higher-order oligomerization. AlphaFold3 predictions produce non-consistent decameric assemblies with similar poor PAE and ipTM scores, showing limited ability to predict a consistent assembly fold. The low signal originating from the multiple sequence alignment (MSA), stemming from the scarcity of homologous sequences, prevented AlphaFold from predicting stable multimeric interactions: none of the multimeric runs (dimer, trimer, tetramer, etc.) yielded PAE or ipTM scores consistent with binding, providing an example of limitation for AlphaFold when sequence alignment depth is insufficient.

**Fig. 3.**
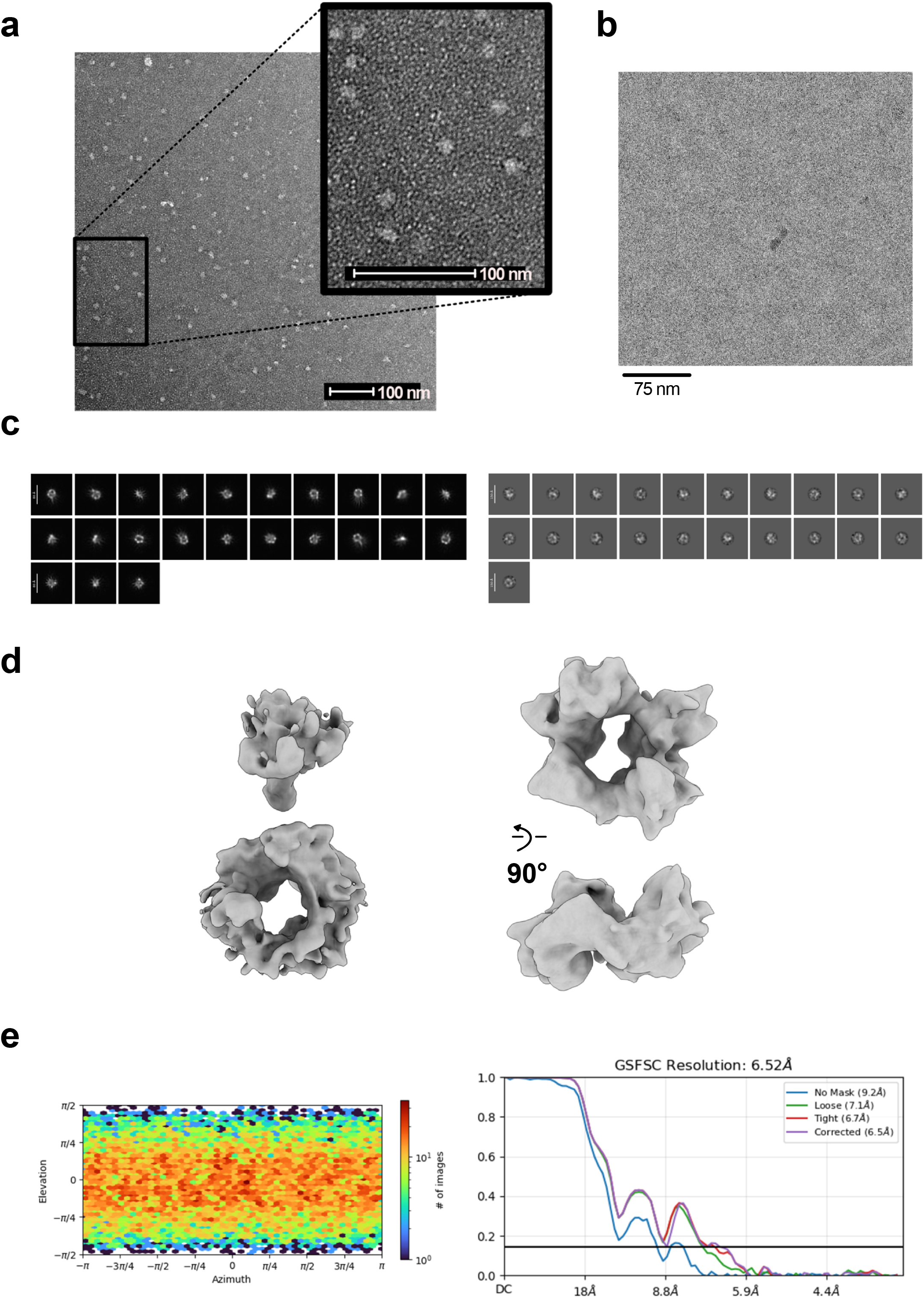
Hydra CryoEM processing. **a**. Negative stain electron microscopy of Hydra. Single particles are observed that are consistent with a decamer. **b**. Representative cryoEM micrograph of the hydra domain. **c**. 2D class averages are shown with non-negativity enforced in order to highlight features and symmetry in the particles (left) and particles used in the reconstruction are shown (right). **d**. Reconstruction of the hydra multimer. The first reconstruction (left) contains an addition feature that is presumed to be 1-2 subunits entering or leaving the decamer. Further rounds of heterogeneous refinement yielded a single ring (on the right) that shows 5 domains. **e**. Angular sampling and gold-standard Fourier Shell Correlation are shown for the final reconstruction.

The negative-stain EM data is not resolved enough to clarify the pseudosymmetric nature of the decamer structures, prompting the use of cryogenic electron microscopy (cryo-EM) to obtain higher-resolution insights into the complex (**Fig. 3b-e**). The resulting 2D classifications supports the idea of a highly dynamic complex, with many different intermediates vitrified on the grid (Fig. 3d). Due to the dynamic nature of the complex, the resolution of the refined maps is limited and lacking a good starting model from AlphaFold, no reliable 3D reconstruction was possible.

### The Hydra domain supports higher-order oligomerization of *Tg*SNF2L

To assess Hydra’s role within the full-length protein, both *Tg*SNF2L and a truncated variant lacking residue 880-1040 (designated *Tg*SNF2LΔhydra) were expressed in baculovirus-infected insect cells, yielding high-purity proteins after purification (**Fig. 4a**). To evaluate the remodeling activity of the recombinant preparations and verify that deletion of the Hydra domain does not compromise catalytic function, we performed a restriction enzyme accessibility assay using EpiDyne nucleosome substrates. In this assay, successful remodeling exposes internal GATC sites, enabling their cleavage by the restriction enzyme DpnII. The efficiency of remodeling was assessed over a time course using an excess of recombinant enzyme (10 nM) to avoid concentration-limited effects. Both *Tg*SNF2L and *Tg*SNF2LΔhydra remodeling activity was benchmarked against commercial human *Hs*SNF2h as a positive control (**Fig. 4b**). All three enzymes catalyzed robust nucleosome remodeling, with similar kinetics and endpoint efficiencies. These results confirm that both recombinant *Tg*SNF2L variants are catalytically active and functionally competent, supporting the structural and mechanistic integrity of the purified proteins.

**Fig. 4.**
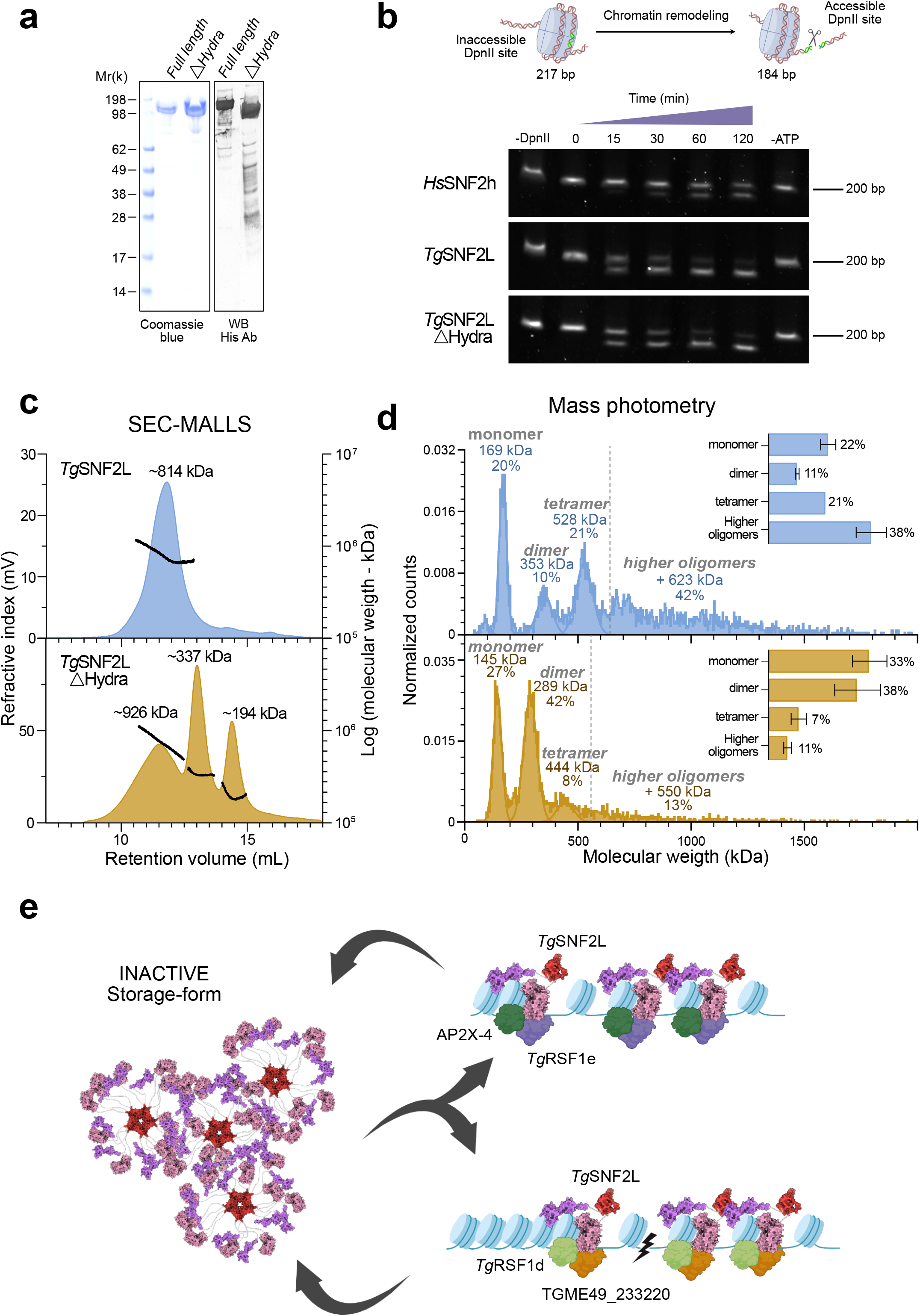
The hydra domain promotes the intrinsic multimerization of *Tg*SNF2L *in vitro*. **a**. Recombinant *Tg*SNF2L and its hydra domain deletion variant (Δhydra) were purified and analyzed by 4-12% NuPAGE, followed by Coomassie blue staining and anti-His tag Western blotting. **b**. Nucleosome remodeling assay using restriction enzyme accessibility confirms that both full-length and Δhydra recombinant *Tg*SNF2L retain catalytic activity. Commercial *Hs*SNF2h (top), recombinant full-length *Tg*SNF2L (middle), and truncated *Tg*SNF2L lacking the Hydra domain (bottom) were incubated with EpiDyne nucleosome remodeling substrates. In this assay, remodeling exposes previously occluded GATC sites, enabling cleavage by the restriction enzyme DpnII. The upper band corresponds to intact nucleosomes; the appearance of the lower band indicates successful remodeling. The first lane serves as a -DpnII control, subsequent lanes represent increasing reaction times and the final lane is - ATP control. **c**. Size-exclusion chromatography coupled with multi-angle light scattering (SEC-MALLS) shows that removing the hydra domain decreases the higher oligomeric forms of *Tg*SNF2L in the micromolar range. With the loss of the hydra domain, two new forms are detected, corresponding to a *Tg*SNF2L and *Tg*SNF2LΔhydra SEC-MALLS (Superose 6 Increase) chromatograms shown as the refractive index curves in blue and orange, respectively. Point measurements of the molecular weight in kDa are displayed as black curves with average masses within the peak regions. **d**. Mass photometry demonstrates a decrease in tetramer and higher oligomeric forms in the nanomolar range upon hydra domain deletion. The data, shown as normalized counts per molecular weight bin (one representative experiment), compares *Tg*SNF2L and *Tg*SNF2LΔhydra in blue and orange, respectively. Monomer, dimer and tetramer peaks are fitted using Gaussian distribution model while higher oligomeric forms are delimited by a dotted line. The relative quantifications of these peaks or windows are shown on the right with the mean and standard deviations shown from duplicate measurements. **e**. Proposed model: The hydra domain acts as a multimerization module, facilitating *Tg*SNF2L storage in a functionally primed state. In this model, *Tg*SNF2L’s multi-oligomeric forms may rapidly release *Tg*SNF2L and its associated proteins in response to DNA damage or replication fork progression.

Having established functional integrity, we next investigated whether the Hydra domain contributes to *Tg*SNF2L oligomerization. Previous analyses demonstrated that the Hydra domain alone forms decameric assemblies *in vitro*, suggesting it may act as a multimerization module. To test whether this property is preserved in the context of full-length *Tg*SNF2L, we used SEC-MALLS to examine oligomerization behavior. *Tg*SNF2L formed higher-order oligomers at high concentrations, with a major peak at approximately 814 kDa, consistent with a tetrameric complex (**Fig. 4c**). Analysis of *Tg*SNF2LΔhydra, concentrated equally prior to injection, showed reduced oligomerization, notably peaks corresponding to dimeric (337 kDa) and monomeric (194 kDa) forms of the protein (**Fig. 4c**). The high micromolar concentration required for a SEC-MALLS analysis is a known factor promoting concentration dependent oligomerization, as such, we also used mass photometry to re-assess this phenomenon within the nanomolar range.

In these conditions, mass photometry provides a more diverse picture of the oligomeric species formed by both *Tg*SNF2L and *Tg*SNF2LΔhydra (**Fig. 4d**). *Tg*SNF2L exists in a multitude of oligomeric states including monomeric, dimeric, tetrameric and above, though discrete sizes beyond the tetramer are not visible. The majority of the particles (59%) are measured as tetrameric and bigger while the dimeric form only represents 11% of the particles and 22% are found as monomers (**Fig. 4d**). The loss of the Hydra domain dramatically affects the *Tg*SNF2L multimerization dynamics *in vitro* (**Fig. 4d**). In a stark contrast to *Tg*SNF2L, monomeric and dimeric forms represent the majority (71%) of the oligomeric states within *Tg*SNF2LΔhydra, with the higher oligomers starting from the tetramer representing only 19% of the remaining species. One notable aspect of this shift is the strong enrichment of dimeric species in *Tg*SNF2LΔhydra (**Fig. 4d**), indicating that hydra does not promote dimerization, which is likely mediated by other domains within the protein. It is also worth noting that trimers were not detected in any sample, suggesting that trimer formation is not an intermediate step toward tetramer assembly. Collectively, these results suggest that the Hydra domain specifically facilitates the formation of higher-order oligomers of *Tg*SNF2L - from tetramer upwards - while disfavoring the formation of stable dimers.

## Discussion

Phylogenetic and structural analyses reveal that *Toxoplasma gondii* evolved two distinct ISWI remodelers, *Tg*SNF2h and *Tg*SNF2L, which preserve the core architecture of their human counterparts but exhibit evolved lineage-specific functional adaptations. The dual presence of these remodelers in coccidian parasites contrasts with their asymmetric loss in other apicomplexans: *Plasmodium* species (hematozoa) have lost SNF2h, whereas piroplasms have lost SNF2L. This pattern supports an ancestral apicomplexan origin of both remodelers, followed by lineage-specific retention reflecting divergent functional needs. A hallmark of *Tg*SNF2L is the Hydra domain - a novel, coccidia-specific insertion absent from all other known ISWI proteins. We previously reported that *Tg*SNF2L assembles into two distinct complexes with non-overlapping functions: one, composed of *Tg*RSF1e and AP2X-4, is essential for parasite viability, while the other, containing *Tg*RSF1d and TGME49_233220, is dispensable to tachyzoites grown *in vitro* (Pachano et al. 2025).

The present findings suggest that the Hydra domain promotes *Tg*SNF2L multimerization. When expressed alone, Hydra self-assembles into a reversible and dynamic equilibrium between monomeric and decameric states. In the context of full-length *Tg*SNF2L, deletion of the domain abolishes higher-order oligomers, favoring the formation of dimers and monomers instead. Interestingly, *Tg*SNF2L-associated complexes purified from *T. gondii* parasites arrested in G1 after egress are homogeneous, with a consistent globular size of ∼1 MDa (Pachano et al. 2025). By contrast, complexes isolated form actively dividing intracellular tachyzoites exhibit pronounced heterogeneity and migrate beyond the upper limit of the size-exclusion column (data not shown). This variability in complex behavior may reflect Hydra-mediated context-dependent multimerization, influenced by cell-cycle or the relative abundance of interacting partners. We propose a model in which Hydra facilitates the formation of ‘reservoir pool’ *Tg*SNF2L complexes that can be mobilized in response to cell-cycle or other cues to assemble and act directly in chromatin-related processes (**Fig. 4e**). While in the initial functional study we did not observe a strong transcriptional effect following the conditional depletion of SNF2L, using the mAID system, recently published results of SNF2L depletion state a more pronounced role in parasite differentiation (Zhu et al. 2025); reconciling these views will require future work.

This broader ability of Hydra to modulate oligomerization prompts comparison with general behavior of nuclear proteins. Although homomultimeric assemblies are common across proteins, nuclear proteins predominantly organize into low-copy discrete oligomers, polydisperse higher-order structures, or phase-separated condensates associated with different nuclear speckles. In the case of *Tg*SNF2L, Hydra - despite behaving as a discrete oligomer when purified alone - drives the full length *Tg*SNF2L protein toward polydisperse oligomers, overriding its intrinsic tendency to form dimers.

The ability of Hydra to drive homo-multimerization may also promote liquid-liquid phase separation (LLPS), transiently sequestering *Tg*SNF2L away from accessible chromatin to fine-tune its availability and activity during nucleosome spacing adjustments following replication or DNA repair. In this model, the *Tg*SNF2L-containing ISWI complexes engage WHIM1 domain proteins (Pachano et al. 2025), which likely act as “protein rulers” to set the distance between adjacent nucleosomes, analogous to their eukaryotic counterparts (Yamada et al. 2011). Overall, the acquisition of Hydra emerges as a key evolutionary hallmark in coccidian ISWI, conferring robust oligomerization capacity and supporting lineage-specific adaptations in chromatin regulation.

## Methods

### Recombinant expression of Hydra

SNF2L Hydra domain (886-1057) was codon optimized for E. coli, synthetized and cloned by Genscript within a modified pET30-a (+) vector (Addgene) in order to possess a C Terminal, TEV cleavable, dual 6*His Tag. Complete sequence can be found in Supplementary Table 1. Expression of the recombinant protein was performed in BL21(DE3) chemically competent cells. Briefly, on day 1, 50 μl of BL21(DE3) cells were incubated with 1 ug of plasmid for 10 minutes at 4°C, transformed by heat shock at 42°C for 45 sec and further incubated 10 min on ice. Following transformation, 600 ul of Luria Broth (LB - Formedium) were added and a 1h, 37°C pre-culture was undertaken before platting 150 ul of pre-culture on LB/Chloramphenicol (Chlo – Sigma Aldrich)/Kanamycin (Kan - Sigma Aldrich) agar plates which were further incubated 12h. On day 2, a single colony was harvested to inoculate 50 ml of LB/Chlo/Kan for 12h at 37°C. On day 3, the 50 ml saturated pre culture then inoculated 3*1L of Terrific Broth/Chlo/Kan (TB - Formedium) expression culture (using a 2 ml inoculum) which was incubated at 37°C. Upon reaching an OD600 of 0.5-0.8, cultures were ice cooled to 20°C for 10 min then induced with 500 μM of IPTG (Euromedex) for 12h after which cultures were centrifuged and stored as dry pellets at - 80°C.

### Hydra purification

For purification, three cell pellets of about 1L of BL21(DE3) culture were each resuspended in 50 ml of lysis buffer [50 mM tris (pH 8.0), 400 mM NaCl, and 2 mM β-mercaptoethanol (BME)] in the presence of an anti-protease cocktail (Complete EDTA-free, Roche). Lysis was performed on ice by sonication for 10 min (30-s on/30-s off, 55° amplitude). Clarification was then performed by centrifugation 1h at 12000g/4°C after which the supernatant was supplemented with 20 mM imidazole and further incubated with 5 ml Ni-NTA resin with a stirring magnet at 4°C for 30 min. Resin retention was performed by gravity with a Bio-Rad glass column after which the resin was washed with 100 mL of washing buffer (50 mM Tris pH: 7.5, 1M NaCl, 2 mM BME and 20 mM Imidazole). His-tagged Hydra was then eluted by in 50 mM Tris pH: 7.5, 300 mM NaCl, 300 mM Imidazole. Size exclusion liquid chromatography was performed on an Akta Purifier in a buffer containing 50 mM Tris pH: 7.5, 150 mM NaCl, 1 mM BME. Following elution, fractions of interest were pooled and concentrated to 0.5 mL using 10 kDa cut-off concentrators before flash frozen in liquid nitrogen and stored at -80°C. Re-injection experiments were also performed using the Akta Purifier system. NuPAGE and NativePAGE gels were run according to the manufacturer’s recommendation.

### Size exclusion chromatography coupled to laser light scattering (SEC-MALLS)

Hydra SEC-MALLS was performed on a S200 10/300 GL increase column (GE Healthcare) running in a buffer system containing 50 mM tris (pH 7.5), 200 mM NaCl, and 1 mM b-ME on an OMNISEC (Malvern) system equipped with RALS, LALS, UV RI and Viscosimeter detectors. Injections of 50 ul were performed using protein samples concentrated at a minimum of 4 mg/ml, and a constant flow rate of 0.5 ml/min was used. Data treatment and mass determination were performed using the OMNISEC software.

### Mass Photometry

Mass photometry measurements were performed using the One MP mass photometer with a concentration of Hydra of 200 nM in 50 mM Tris (pH 7.5), 200 mM NaCl, 1 mM BME. Importantly, the blank buffer acquisition was performed on buffer filtrated through an Amicon ultra 30 kDa concentrator. Standard protein mass measurements were also performed within the same buffer system.

### Protein structure predictions

AlphaFold3 (Abranson et al., Nature 2024) predictions were run on the google AlphaFold server. All models were depicted using UCSF ChimeraX (Pettersen et al. 2020).

### Negative stain electron microscopy

Purified Hydra domain samples were thawed and diluted for application on carbon coated copper grids (300 mesh, Electron Microscopy Science). 2 % uranyl acetate solution was used for staining and grids left to dry before loading on a FEI Tecnai T12 electron microscope operating at 120 kV and equipped with a Ceta 16 M camera. Data was manually collected at magnifications between 30kx and 68kx. Images were stored as TIFF files.

### Cryo-electron microscopy

Purified Hydra domain samples were thawed and diluted for application on carbon 1.2/1.3 Cu 300 Quantifoil grids (Electron Microscopy Science). Prior to sample application, the grids were hydrophilized with a NanoClean 1070 plasma cleaner (Fischione) at 100% power, 25:72 oxygen:argon gas mix for 1:30 min. Grid vitrification was performed on a Vitrobot Mark IV (FEI), applying 3.5 µl sample per grid, blotting for 4 s at blot force -1. Grids were then clipped and data collected on a FEI Talos Glacios electron microscope operating at 200 kV (EMBL Grenoble). Micrographs were collected using EPU from a Falcon 4i direct electron detector with a SelectrisX energy filter. A total of 3800 movies of 1053 EER frames were collected at 130 000 Magnification, a pixel size of 0.878 Å with a total exposure of 40 e-/Å2. The movies were imported into the cryoSPARC software suite. Movies were aligned, motion-corrected and CTF estimated prior to particle picking. For initial picking, low resolution templates generated from negative stain data were used, followed by iterative Topaz training to yield particles of better quality. Multiple rounds of 2D classifications were run for particle alignment. The particles were heterogeneous and had a high background but 2D class averages. Rounds of homogenous refinement yielded a final map of the sample at low resolution showing the ring structure with an additional feature next to the ring. As the assembly of hydra is dynamic, we presume that the extra feature is one or two subunits leaving or entering the ring. Further rounds of heterogeneous refinement allowed the selection of particles containing just the ring structure.

### Insect cell recombinant expression and purification *Tg*SNF2L and *Tg*SNF2LΔHydra

Codon optimized *Tg*SNF2L (aa 1 to 1556) and *Tg*SNF2LΔHydra (aa 1 to 1556 with an internal deletion between aa 880 and 1040) were synthetised by Genscript and inserted within the pFastBac vector. The baculovirus generation and insect culture protocol is identical to previous work (Robert et al. 2025).

### Nucleosome remodeling assay

Nucleosome remodeling reactions were performed using the EpiDyne Nucleosome Remodeling Assay Substrate ST601-GATC1 (EpiCypher, SKU: 16-4101). Recombinant full-length TgSNF2L and TgSNF2LΔHydra were used alongside human SNF2h (HsSNF2h/SMARCA5, EpiCypher, SKU: 15-1024) as a positive control. Reactions were assembled in 10 µL final volume in Remodeling buffer (20 mM Tris pH 7.5, 50 mM KCl, 3 mM MgCl□, 0.01% Tween-20, 0.01% BSA). Reaction components were added sequentially in the following order: 10 nM remodeler enzyme (2.5 µL), 20 nM nucleosome substrate (2.5 µL), 10 units DpnII (New England Biolabs, R0543S; 2.5 µL), and 2 mM ATP (2.5 µL). Reactions were incubated at 30°C, and aliquots were removed at designated time points for kinetic analysis. Control reactions (no DpnII, no ATP) were stopped after 120 minutes. Reactions were terminated by adding an equal volume of Quenching buffer (10 mM Tris pH 7.5, 400 mM EDTA, 0.6% SDS, 50 µg/mL Proteinase K), followed by incubation at 55°C for 20 min. Reaction products were resolved by 6% TBE-Urea gel electrophoresis (run at 12 W constant power), stained with SYBR Safe for 10minutes (Thermo Fisher Scientific), and visualized by an Ibright CL750 (Invitrogen) fluorescence imaging.

### Statistics and Reproducibility

Sample sizes were not predetermined and chosen according to previous literature. Experiments were performed in biological replicates and provided consistent statistically relevant results. No method of randomization was used. All experiments were performed in independent biological replicates as stated for each experiment in the manuscript.

## Data availability

Correspondence and requests for materials should be addressed to C.S.

## Acknowledgments

We are grateful to the developers of the ToxoDB.org Genome Resource. ToxoDB and EuPathDB are part of the National Institutes of Health/National Institutes of Allergy and Infectious Diseases (NIH/NIAID)-funded Bioinformatics Resource Center. This work was supported by MSD Avenir [Project LatentToxoDiag, DS-2022-0017], the Laboratoire d’Excellence (LabEx) ParaFrap [ANR-11-LABX-0024], Fondation pour la Recherche Médicale [FRM Equipe, EQU202103012571] and the Agence Nationale pour la Recherche [Project ApiNewDrug, ANR-21-CE35-0010-01; Project ApiMORCing, ANR-21-CE15-0002-01; Project Chaires Excellence Biologie Santé_ToxoNeoSex, ANR-24-CHBS-0008]. We would like to thank Iskander Khusainov and Romain Linares for excellent support in using the EM Facility at EMBL Grenoble. This work used the platforms of the Grenoble Instruct-ERIC center (ISBG; UAR 3518 CNRS-CEA-UGA-EMBL) within the Grenoble Partnership for Structural Biology (PSB), supported by FRISBI (ANR-10-INBS-0005-02) and GRAL, financed within the University Grenoble Alpes graduate school (Ecoles Universitaires de Recherche) CBH-EUR-GS (ANR-17-EURE-0003).

## Author Contributions

C.S. and M.W.B supervised the research and coordinated the collaboration. B.P., J.V., C.C., C.M., L.P., M.-A.H, M.W.B, and C.S. designed, performed and interpreted the experimental work. B. P, M.-A.H. and C. S. wrote the paper with comments from all other authors.

## Declaration of Interests

The authors declare no competing interests.

## Supplementary data

**Supplementary Figure 1** | **Superose 6 separation of Hydra. a**. Re-injection of a previously purified monomeric fraction of hydra on a superose 6 10/300 GL increase column running in 30mM Tris pH7.5, 150 mM NaCl, 5 mM BME. **b**. NuPAGE gel colored with Coomassie blue of the separated fractions. **c**. NativePAGE gel colored with Coomassie blue of the separated fractions.

**Supplementary Table 1** | **Description of molecular biology reagents**. DNA synthesis constructs used in this work are charted in the table.

## Notes

### Competing Interest Statement

The authors have declared no competing interest.

