## Supplementary figures and images for "Hydra domain drives SNF2L multimerization and marks ISWI diversification in parasites"

### Supplementary Figure 1

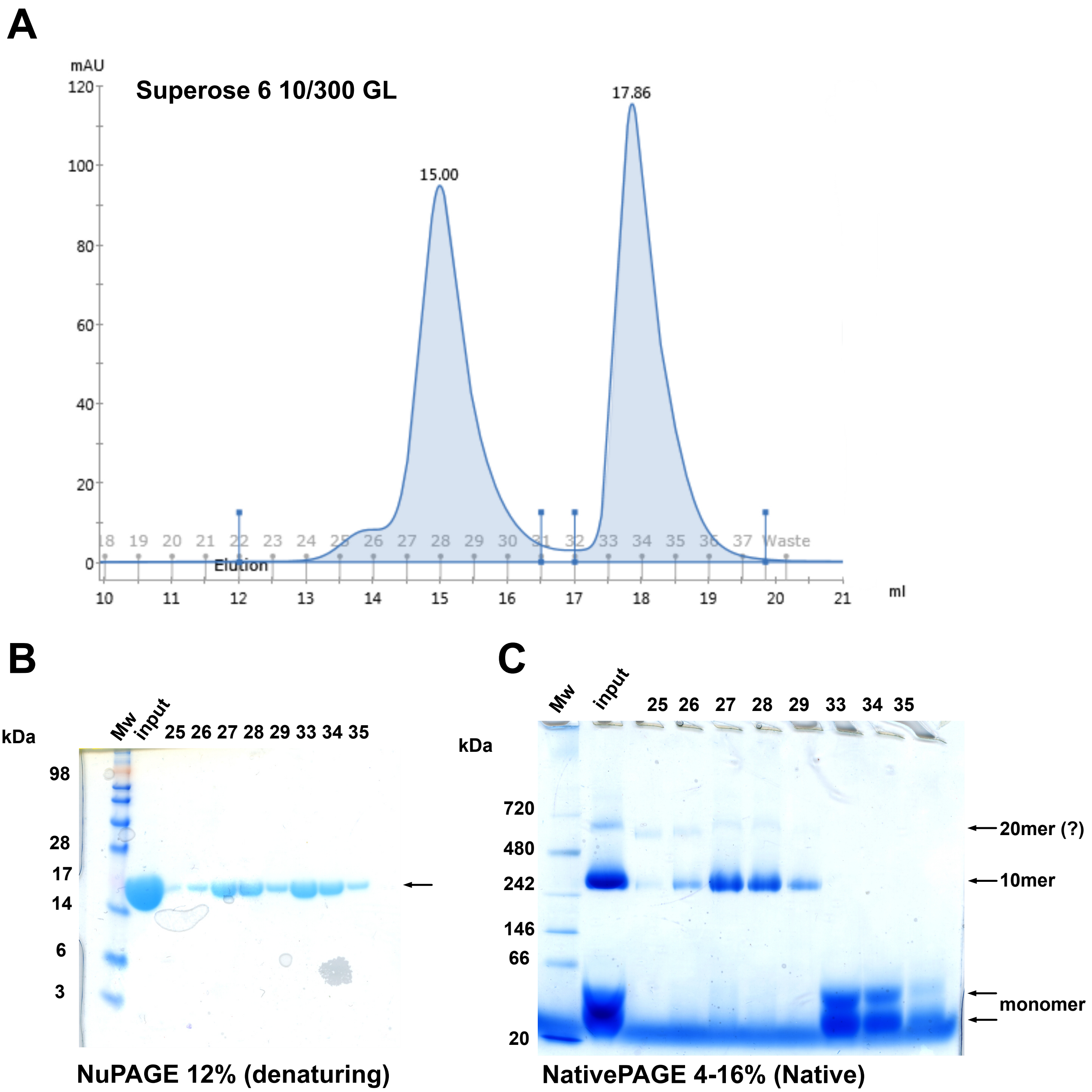
